# DNA repair under heat: DNA Polymerase λ modulates heat stress-induced mutagenesis in plants

**DOI:** 10.1101/2025.04.24.650460

**Authors:** Clair M. Wootan, John Lutterman, Nathan Springer, Xiaosa Xu, Feng Zhang

## Abstract

Mutation rates can increase substantially under environmental stress, known as stress-induced mutagenesis. Specifically, heat stress has been shown to elevate mutation rates, thereby enhancing genetic variability and facilitating adaptation. However, the underlying mechanisms remain elusive in eukaryotes. Here, we investigated how heat stress affects DNA repair and induces mutations both locally and globally through CRISPR-Cas9 targeted DNA breaks and whole genome sequencing analyses in *Arabidopsis thaliana*. Heat stress was found to enhance CRISPR editing efficiency across all chromatin contexts, with particularly significant increases, up to 29.9-fold, in heterochromatic regions. Moreover, heat stress consistently shifts mutation outcomes toward 1-bp insertions regardless of chromatin states. We identified a heat-inducible, error-prone DNA polymerase, Polλ, as the key mediator of mutation profile changes. When extending our investigation from targeted mutations to genome-wide effects, we found that increases in global mutation rates under heat stress are also dependent on Polλ. Single-cell transcriptomic analysis further demonstrated that Polλ expression is tightly regulated and cell-type specific, with the highest expression levels in central zone meristematic cells. Together, these findings provide practical applications for improving editing efficiency in heterochromatic regions and fundamental insights into heat-induced mutagenesis, establishing Polλ as a crucial mediator of stress-induced genetic variation in plants.

## Main

Environmental stress can increase mutation rates, a phenomenon known as stress-induced mutagenesis (SIM), enhancing evolutionary potential and adaptive flexibility ^1,2^. Among diverse environmental stressors, heat stress has emerged as a significant abiotic threat facing organisms worldwide ^3^. As global temperatures continue to rise due to climate change, understanding the genomic and molecular impacts of heat stress is becoming increasingly critical for both fundamental biology and applied sciences ^4,5^. Elevated temperatures can increase mutation rates in both prokaryotic and eukaryotic organisms ^5^. The increased mutation rates are thought to be the results from two primary factors, direct induction of DNA lesions and upregulation of error-prone DNA repair machinery under heat stress. Heat stress can directly induce to various forms of DNA damages, including the oxidative lesions (e.g., 8-oxoguanine, deaminated cytosine, and apurinic sites), single-stranded DNA breaks (SSBs), and double-stranded DNA breaks (DSBs) ^6^. In contrast to these well-documented DNA lesions, the mechanisms by which heat stress alters DNA repair pathways remain less characterized except for in bacteria ^7,8^.

Plants, due to their sessile nature and constant exposure to environmental fluctuations, provide a valuable system for investigating the genomic consequences of thermal stress. Genome-wide analyses in *Arabidopsis thaliana* have revealed heat-induced increases in mutation rates and altered mutational spectra ^9,10^. More recently, a heat-responsive module involving heat shock protein 90 (*HSP90*), high expression of osmotically responsive genes (*HOS1*), and the RecQ helicase L2 (*RECQ2*) was found to play a critical role in maintaining DNA integrity under heat stress, beginning to identify direct links between heat stress and DNA repair regulation in plants ^11^. Despite these advances, the molecular mechanisms underlying heat-induced genomic mutation rate increase remain largely unknown in plants. In this study, we sought to address how heat stress affects DNA repair and induces mutations across diverse chromatin contexts through systematic investigation of CRISPR-Cas9 targeted DNA breaks and whole genome sequencing analyses in *Arabidopsis*.

### Heat stress significantly increases CRISPR editing efficiency across chromatin contexts

We first employed a targeted DNA damage approach using CRISPR-Cas9. As outlined in Figure 1A, we applied a previously established heat stress protocol to *Arabidopsis* seeds, exposing them to 37°C for 24 hours during germination, followed by a 24-hour recovery period at 22°C ^12^. This heat stress and recovery cycle was repeated up to three times. Following the treatment, seedlings were grown under long-day conditions at 22°C for two weeks prior to tissue collection, DNA extraction, and Next-Generation Sequencing (NGS) to assess gene editing outcomes. To determine the effects of heat stress on CRISPR-mediated editing efficiency across chromatin contexts, we utilized a previously developed *Arabidopsis* line that stably expresses a CRISPR-Cas9 T-DNA construct targeting a multi-copy CRISPR target site (MC4 site) ^13^. This construct includes two guide RNAs (gRNAs): one targeting the endogenous *Chetelase I 2* (CHLI2) gene and the other targeting a 20-bp sequence repeated at eight unlinked loci with varying chromatin environments. Based on previously characterized epigenetic features, these loci were classified into two groups: high-accessibility and low-accessibility sites.

**Figure 1.**
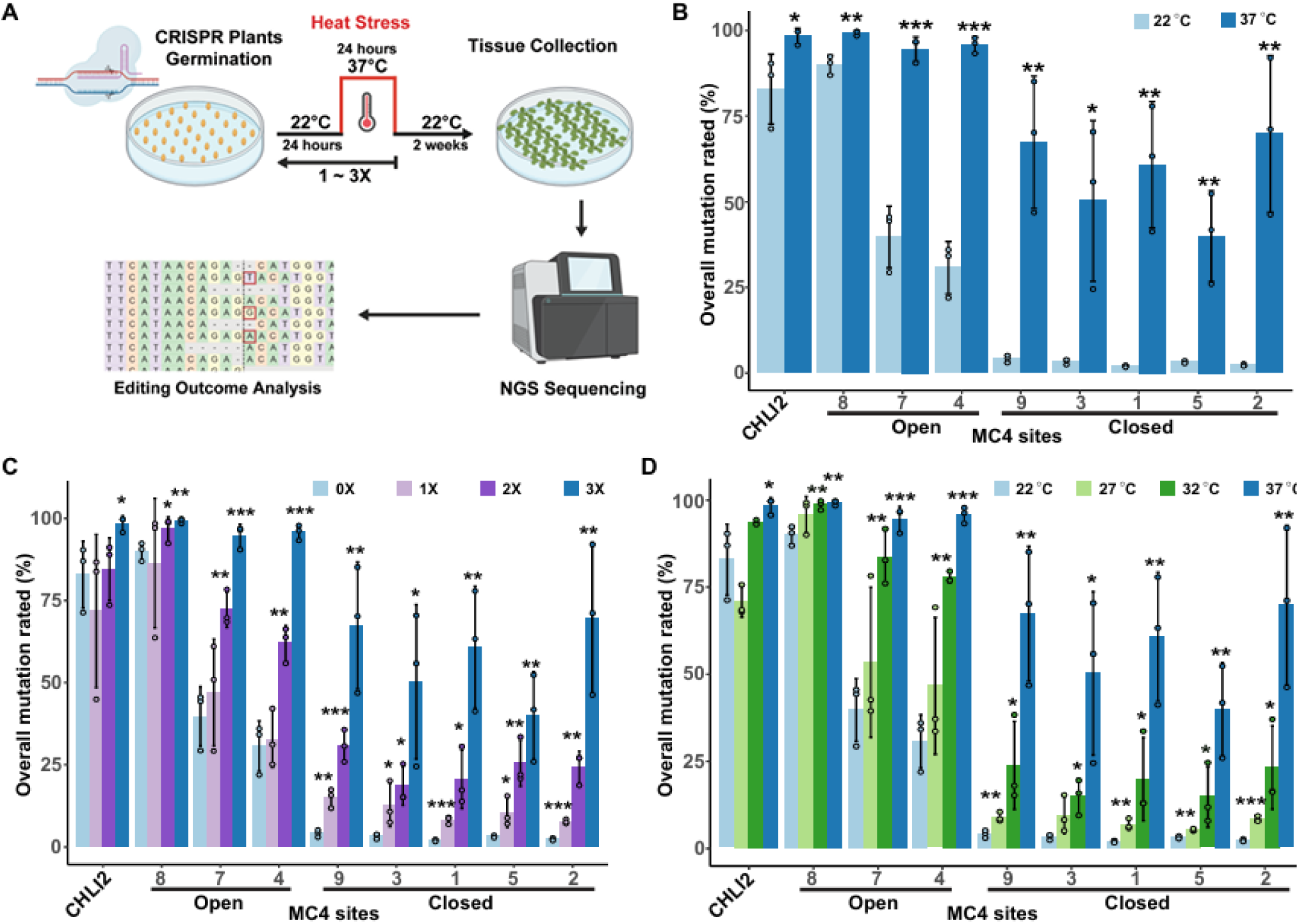
Heat stress increases editing efficiency across diverse chromatin landscapes. **a,** Schematic workflow of the heat stress protocol, showing elevated temperature treatments applied for 1-3 cycles during seed germination, followed by a 2-week recovery period at standard growth conditions. **b,** Editing efficiencies for CHLI2 and individual MC4 sites. Light blue represents control conditions and dark blue indicates heat stress treatment. **c,** Editing efficiencies with duration treatments at 0x (light blue), 1x (light purple), 2x (dark purple), and 3x (dark blue) heat stress cycles at 37°C. **d,** Editing efficiencies with temperature gradient treatments at 27°C (light green), 32°C (dark green), and 37°C (dark blue) with 3x heat stress cycles. The *p*-values were calculated with t-tests comparing each heat stress sample to the corresponding control for each site and treatment (n=3; \**p* < 0.05; \*\**p* < 0.005, \*\*\**p* < 0.0005 ). The source data are provided in the Source Data file.

Under control conditions, the CHLI2 target site exhibited an average CRISPR editing efficiency of 82.8% (Fig. 1B). After heat stress, CHLI2 editing efficiency significantly increased to 98.2%, consistent with previous findings that gene editing efficiency is typically higher in genic regions. We next analyzed editing efficiencies at the eight multi-copy sites distributed across chromatin contexts. In agreement with prior findings, a strong correlation between chromatin accessibility and editing efficiency was observed under control conditions: high-accessibility sites displayed editing efficiencies ranging from 30.7% to 89.9%, while low-accessibility sites exhibited lower efficiencies, ranging from 2.0% to 4.1%^13^ (Fig. 1B).

Under the heat stress condition, editing efficiencies were enhanced across nearly all multi-copy target sites, irrespective of chromatin context. This widespread improvement was evident in both euchromatic and heterochromatic sites, with a single exception at Site 8, where editing efficiency remained similar, likely due to saturation at its maximum level (Fig. 1B). In euchromatic regions, editing efficiencies increased moderately yet significantly, rising 1.1- to 3.1 - fold at Sites 4 and 7, respectively. Notably, heterochromatic regions exhibited dramatically higher enhancement in response to heat treatment, with significant fold increases from 12.8 to 29.9-fold at Sites 9, 3, 1, 5, and 2, respectively (Fig. 1B).

As previous studies suggested positive correlations between temperature and CRISPR-Cas9 editing efficiencies in euchromatic regions ^12,14^, we examined how varying both temperature and duration affected editing efficiencies across CHLI2 and the MC4 sites. When varying stress durations, we exposed germinating seeds to one, two, or three cycles of 24-hour heat stress at 37°C. All sites except the saturated Site 8 and CHLI2 showed increasing editing efficiency with each additional cycle (Fig. 1C). This effect was particularly pronounced at low-accessibility sites, where editing increased 3.2 to 4.0-fold after one cycle, 5.5 to 10.3-fold after two cycles, and 12.3 to 29.9-fold after three cycles. To evaluate temperature sensitivity, we subjected germinating seeds to three cycles of heat stress at 27°C, 32°C, or 37°C. Notably, editing efficiency increased at all unsaturated sites, with stronger effects in low-accessibility regions (Fig. 1D). The low-accessibility sites showed fold increases of up to 3.6 fold at 27°C, up to 9.9 fold at 32°C, to up to 29.9 fold at 37°C. Collectively, these findings demonstrate that heat stress significantly enhances CRISPR-Cas9 editing efficiency across diverse chromatin contexts, with particularly pronounced effects at low-accessibility sites, while both higher temperatures and longer exposures progressively produce stronger effects.

### Heat stress alters CRISPR mutation outcomes toward 1-bp insertions

We next investigated how heat stress influences DNA repair outcomes. Cas9 generates both blunt and staggered cuts three or four nucleotides upstream from the Protospacer Adjacent Motif (PAM) ^15^. Subsequent repair processes lead to mutations that typically fall into three categories: single base insertions, small deletions (1–10 bp), and large deletions (>10 bp) (Fig. 2A). Among these, single base insertions and small deletions account for more than 80% of all mutations (Supplemental Fig. 1A). The ratio between different types of insertions/deletions (indels) reflects distinct DSB repair processes ^15^, therefore providing insight into the impact of heat stress on DNA repair.

**Figure 2:**
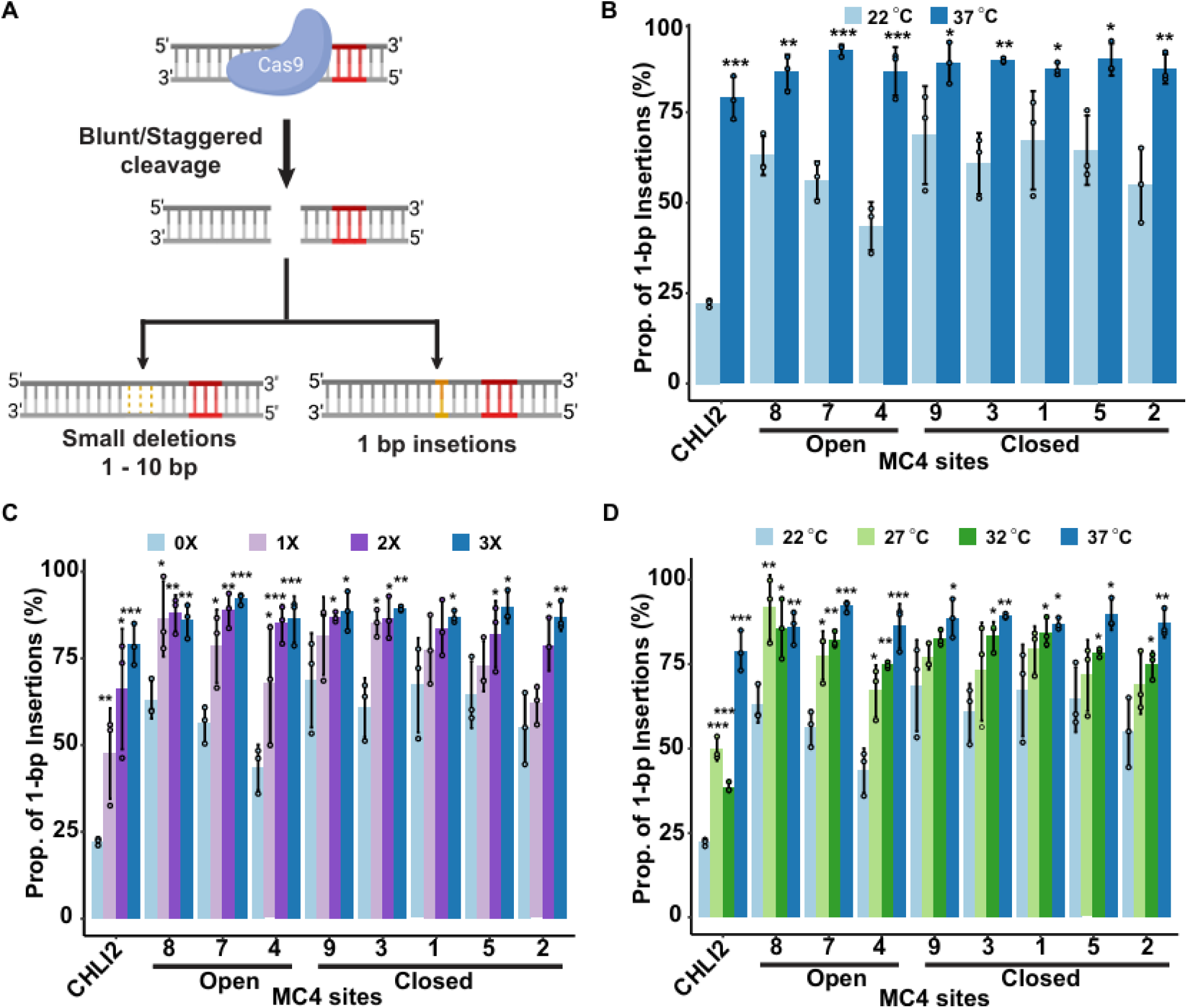
Heat stress alters DNA repair profiles. **a,** Schematic of CRISPR-Cas9 mediated double-stranded DNA repair. **b**, Proportion of 1-bp insertions for CHLI2 and individual MC4 sites. Light blue indicates control conditions and dark blue represents the heat stress treatment. **c,** Proportion of 1-bp insertions with duration treatments at 0x (light blue), 1x (light purple), 2x (dark purple), and 3x (dark blue) heat stress cycles at 37°C. **d**, Proportion of 1-bp insertions with temperature gradient treatments at 22°C (light blue) 27°C (light green), 32°C (dark green), and 37°C (dark blue) with 3x heat stress cycles. The *p*-values were calculated with t-tests comparing each heat stress sample to the corresponding control for each site and treatment (n=3; \**p* < 0.05; \*\**p* < 0.005; \*\*\**p* < 0.0005). The source data are provided in the Source Data file.

When examining repair outcomes under control conditions, we observed distinct indel patterns at the CHLI2 and MC4 sites with varied frequencies of 1-bp insertions. The CHLI2 site exhibited 1-bp insertions at 25% and small deletions at 62%; at MC4, the frequency of 1-bp insertions ranged from 45% to 70%, with small deletions at 26% to 47% (Fig. 2B; Supplemental Fig. 1B). After exposing plants to three cycles of 37°C heat stress, both CHLI2 and MC4 sites displayed significantly increased 1-bp insertions by 1.2 to 3.6 fold (Fig. 2B; Supplemental Fig. 2).

To characterize the sensitivity of this heat-induced repair shift, we varied both temperature and exposure duration as described above. Our duration experiments revealed positive correlations: a single 24-hour heat stress cycle significantly increased 1-bp insertion rates at CHLI2 and five of eight MC4 sites, with all sites showing significant increases after two cycles (Fig. 2C). The magnitude of this effect continued to increase with the third cycle, suggesting a cumulative response to prolonged heat exposure. Similarly, our temperature gradient experiments revealed changes in mutation outcomes even at moderate temperatures, with 27°C and 32°C treatments significantly increasing 1-bp insertion rates at CHLI2 and four of eight MC4 sites, with the 37°C treatment producing the strongest effect overall (Figure 2D).

Notably, both temperature and duration effects on insertion rates displayed chromatin context dependence. While editing efficiency increased more dramatically in low-accessibility sites, the 1-bp insertion rates showed greater sensitivity to heat stress in high-accessibility regions. Specifically, euchromatic regions displayed significant increases in 1-bp insertion rates (1.3- to 2.4-fold) at temperatures as low as 27°C and following just one cycle of heat exposure (1.4- to 2.1-fold). In contrast, heterochromatic regions required both higher temperatures and extended exposure durations to achieve significant changes in 1-bp insertion frequencies (Fig. 2C and D). This chromatin dependent response suggests that chromatin accessibility could play a critical role in modulating the DNA repair processes.

### Heat-inducible DNA Polλ is responsible for increased 1-bp insertion rates

The observed increase in 1-bp insertion rates under heat stress prompted us to investigate the underlying molecular mechanism. Recent work has identified DNA polymerase λ (DNA Polλ) as responsible for CRISPR-mediated 1-bp insertions in *Arabidopsis* ^15^. DNA Polλ is a member of the X family polymerases and is the only homolog in plants, which through its lyase and PolX domains are involved in gap-filling for a range of different repair pathways ^16^.

We then examined whether DNA Polλ is involved in heat-induced 1-bp insertions. Since DNA repair mutants could display compromised heat tolerance, as reported for the *hos1* mutants ^11^, we first assessed heat sensitivity of the *atPolλ-1* mutant plants. We evaluated seedling survival rates and root elongation under heat stress, using wild-type and *hos1* mutant plants as controls. Following our established 3× 37°C heat stress protocol, the *hos1* mutant displayed severe survival defects (10% survival) consistent with the previous report. In contrast, the *atPolλ-1* plants showed no significant heat sensitivity compared to wild-type plants with comparable survival rates (wildtype: 89%; *atPolλ-1*: 85%) (Fig. 3B; Supplemental Fig. 3). Root length comparisons between wildtype and *atPolλ-1* also showed no significant differences during control (wildtype: 0.84 inches; *atPolλ-1:* 0.89 inches) or post heat stress (wildtype: 0.34 inches; *atPolλ-1:* 0.30 inches) (Fig. 3C). These results indicate that DNA Polλ deficiency does not significantly affect overall heat tolerance in *Arabidopsis*.

**Figure 3:**
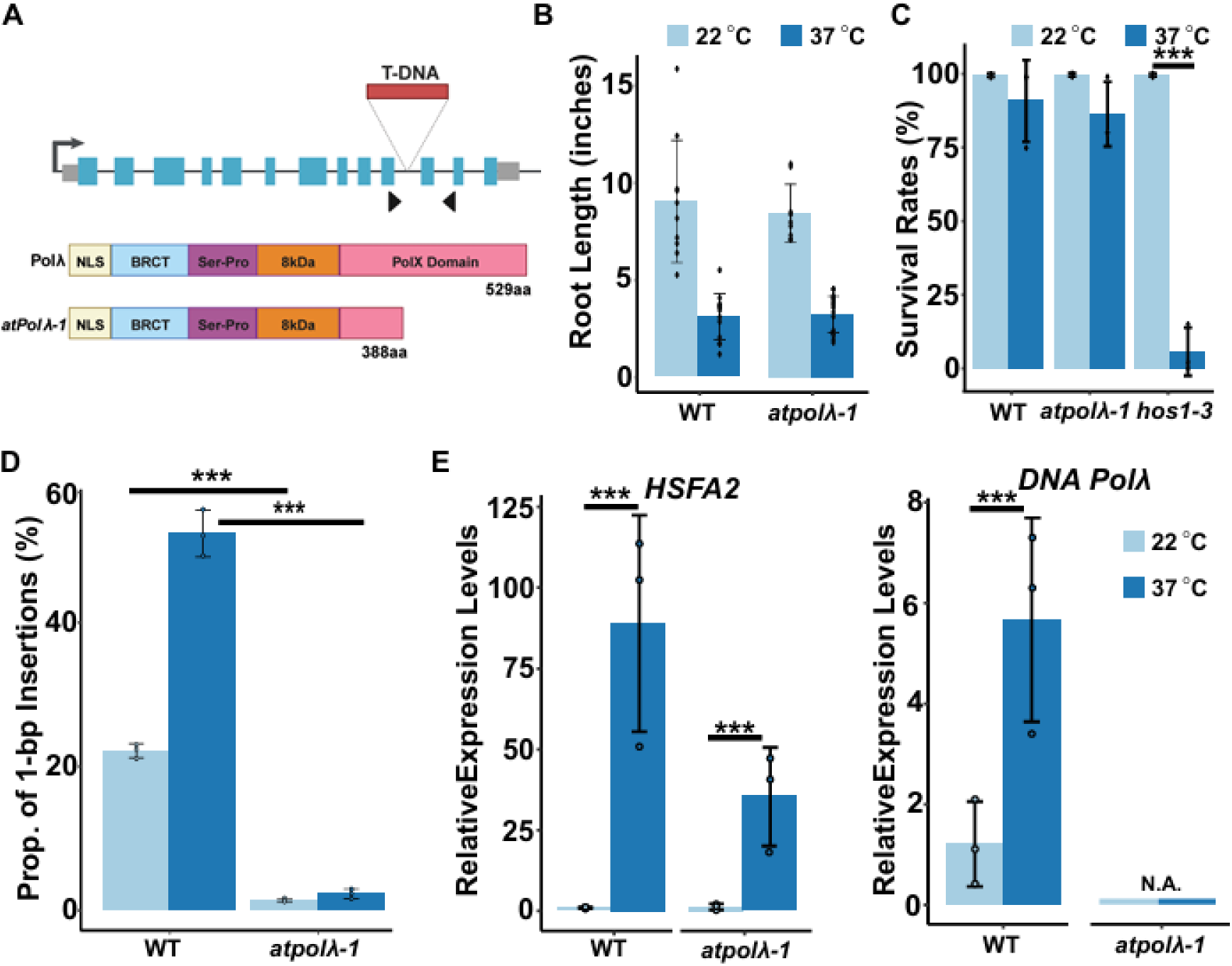
DNA Polλ is heat responsive and responsible for altered repair outcomes under heat stress. **a,** Schematic map of the T-DNA insertion in the *atPolλ-1* mutant. Key protein domains of DNA Polλ include: Breast Cancer Carboxy-terminal (BRCT), Serine/Proline-rich (Ser/Pro), 8kDa, and Catalytic Core or Polymerase domain (PolX). The *atPolλ-1* mutant protein contains a truncated PolX domain. **b,** Root length comparisons between control and heat stress conditions from wildtype and the *atPolλ-1* mutant plants (n = 9 - 18). **c**, Comparisons of seedling survival rates at 37°C for 3 days from wildtype and the *atPolλ-1* and *hos1* mutant plants (n= 33-66). **d**, Comparisons of 1-bp insertion rates between wild type and *atPolλ-1* mutant plants at the CHLI2 sites under control and heat stress conditions. **e,** Comparisons of DNA Polλ expression levels in wildtype and *atPolλ-1* mutant under control and heat stress conditions. The relative expression levels of DNA Polλ were normalized to the housekeeping gene, EF1α. The heat-inducible gene, HSFA2, was included as the positive control. The *p*-values were calculated with t-tests comparing each heat stress sample to the corresponding control for each site and treatment (n=3; \**p* < 0.05, \*\**p* < 0.005, \*\*\**p* < 0.0005 ). The source data are provided in the Source Data file.

To test DNA Polλ’s role in heat-induced 1-bp insertion bias, we generated stable transgenic lines carrying CRISPR-Cas9 constructs targeting CHLI2 in the wild-type and *atPolλ-1* backgrounds. T2 plants were subjected to our 37°C heat stress protocol. Mutation analysis at the CHLI2 site revealed that, while wild-type plants showed the characteristic heat-induced increase in 1-bp insertions, the *atPolλ-1* plants consistently maintained very low levels of 1-bp insertions regardless of temperature treatment (Fig. 3D). These findings indicated that DNA Polλ is crucial for the heat-induced shift toward increased 1-bp insertions.

Previous studies have suggested that DNA Polλ expression can be stress-inducible ^16^. This led us to investigate whether heat could induce expression of DNA Polλ. Quantitative RT-PCR (RT-qPCR) was performed on CRISPR-containing plants in both wild-type and *atPolλ-1* backgrounds directly following the 3× heat stress treatment. We included the well-characterized Heat Shock Transcription Factor A2 (HSFA2) as a positive control to verify effective heat stress treatment ^17,18^. Under control conditions at 22°C, both DNA Polλ and HSFA2 showed low relative gene expression (1.2 and 1.02 in 2|^-ΔΔCt, respectively) as compared to the ELFa1 housekeeping gene control (Fig. 3E). Following heat stress treatment, HSFA2 expression increased dramatically (86-fold in wildtype and 28-fold in *atPolλ-1*), confirming the effectiveness of our heat treatment. Importantly, *DNA Polλ* expression showed a significant 4.6-fold increase in the wild-type background while remaining at undetectable levels in the *atPolλ-1* mutant (Fig. 3E). Taken together, these results demonstrated heat stress-inducible expression of DNA Polλ, which in turn could be involved in shifting the DNA repair outcomes towards 1bp insertions at CRISPR-Cas9-induced double-stranded breaks.

### DNA Polλ modulates genome-wide mutagenesis under heat stress

Given DNA Polλ’s diverse role in different DNA repair pathways and its lack of proofreading abilities, we hypothesized that it could be a key contributor to increased genome-wide mutation rates under heat stress conditions ^19,20^. We then conducted whole genome sequencing (WGS) to assess mutation rates in wild-type and *atPolλ-1* mutant plants under both control and heat stress conditions (Fig. 4A). Following the established computational pipeline for identifying genome-wide *de novo* mutations ^21^, we performed WGS on 3-week-old rosette leaves from wild-type and *atPolλ-1* plants, which were grown under either control conditions or subjected to our 3× heat treatment protocol (Fig. 4A). Three biological replicates were analyzed for each genotype and treatment combination, with sequencing coverage ranging from 85 to 170-fold depth across individual samples.

**Figure 4:**
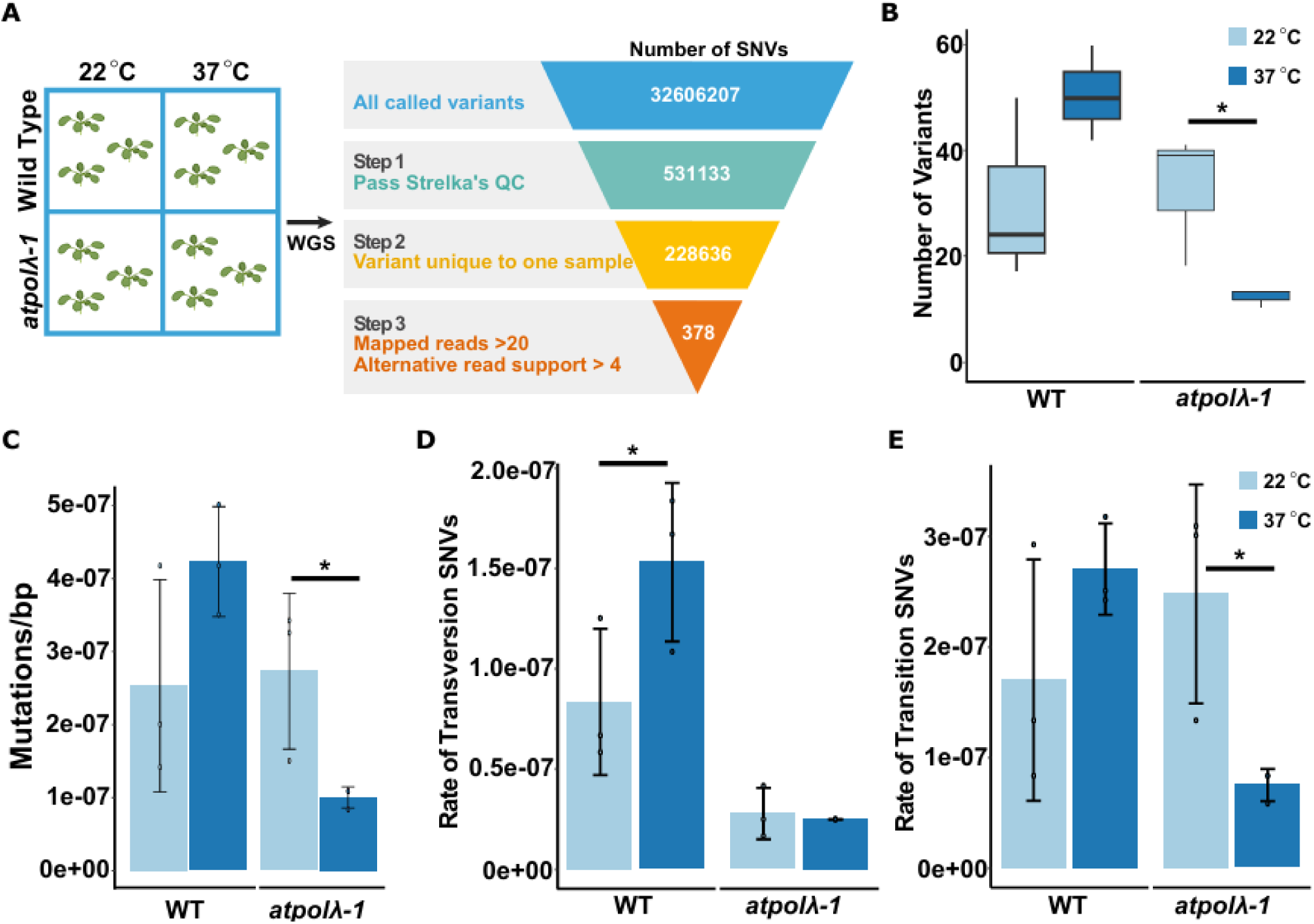
DNA Polλ contributes to genome-wide mutagenesis under heat stress. **a,** Schematic workflow of the whole genome sequencing (WGS) to identify *de novo* mutations from wild type and the *atPolλ-1* mutant plants under control and heat stress conditions. **b**, Comparisons of total number of *de novo* SNVs per genotype under heat stress and control conditions. **c**, Comparisons of mutation rates (total number of SNVs per base pair) between wild type and *atPolλ-1* mutant under control and heat stress conditions. **d**, Comparisons of transversion mutation rates between wild type and *atPolλ-1* mutant under control and heat stress conditions. **e**, Comparisons of transition mutation rates between wild type and *atPolλ-1* mutant under control and heat stress conditions. The *p*-values were calculated with t-tests comparing each heat stress sample to the corresponding control for each genotype, for each treatment (n=3; \**p* < 0.05). The source data are provided in the Source Data file.

To identify *de novo* mutations, we used the variant caller Strelka2 in somatic TUMOR-NORMAL mode, followed by a stringent filtering process to minimize false positives (see Methods; Figure 4A). A high-confidence set of *de novo* 143 small indels (from 3 bp deletion to 3 bp insertions) and 378 single nucleotide variants (SNVs) was obtained across all samples (Fig. 4B, Supplemental Fig. 4A). Our analysis revealed that heat stress did not significantly increase the number of indels in wild-type plants compared to control (14 vs. 12 indels/plant on average, 1.0 × 10⁻⁷ vs. 1.22 × 10⁻⁷ indel rate) while *atPolλ-1* plants under heat stress conditions had significantly fewer small indels compared to *atPolλ-1* control (6 vs. 14 indels/plant on average,1.1 × 10⁻⁷ vs. 5.0 × 10⁻⁸ indel rate; Supplemental Fig. 4B). This suggests that DNA Polλ could be an important contributor to indel mutations under heat stress.

Interestingly, we found heat stress substantially increased the number of *de novo* SNVs in wild-type plants compared to control conditions (50 vs. 30 mutations/plant on average; Fig. 4B). In contrast, *atPolλ-1* plants exhibited fewer mutations under heat stress compared to control conditions (12 vs. 32 mutations/plant on average; Fig. 4B). When normalized to genomic size, the mutation rates for both wild-type and *atPolλ-1* plants under control conditions were comparable (2.2 × 10⁻⁷ and 2.4 × 10⁻⁷ mutations per base pair, respectively), indicating that the absence of DNA Polλ does not alter background mutation rates under normal growth conditions (Fig. 4C). Under heat stress, however, wild-type plants showed elevated mutation rates (3.7 × 10⁻⁷ mutations per base pair; Fig. 4C), while *atPolλ-1* plants exhibited reduced mutation rates (8.8 × 10⁻⁸ mutations per base pair; Fig. 4C). These observations confirm that heat stress substantially increases genome-wide mutation (SNV) rates in *Arabidopsis*. More importantly, DNA Polλ plays a critical role in mediating heat-induced mutagenesis.

Previous studies have demonstrated that DNA Polλ repairs oxidized guanine (8-oxoG) lesions in plants with a characteristic bias toward transversion mutations (C:G→A:T) ^19^. To further investigate the role of DNA Polλ in heat-induced mutagenesis, we categorized the *de novo* SNVs into transitions (C↔T or G↔A) and transversions (C↔A or G↔T) (Supplemental Fig. 4C). In control conditions, both genotypes showed predominantly transition mutations, but wild-type plants displayed a substantially higher proportion of transversions compared to *atPolλ-1* plants (transversion ratio of ∼0.5:1 in wild-type versus ∼0.1:1 in atPolλ-1 plants; Fig. 4D and E). This 5-fold difference confirms DNA Polλ’s role in generating transversion mutations. Under heat stress, wild-type plants showed proportional increases in both transition and transversion mutations, maintaining the ∼0.5:1 ratio (Fig. 4D and E). Notably, *atPolλ-1* plants exhibited a different pattern: while transversion rates remained similar to control conditions, transition mutations decreased dramatically by ∼3.2-fold (from 2.4 × 10⁻⁷ to 7.5 × 10⁻⁸ mutations per base; Fig. 4D and E). These data suggest that DNA Polλ not only facilitates transversions but could also be involved in promoting transition mutations during heat stress. Together, our WGS analyses suggest the strong contribution of DNA Polλ to mutation rate changes under heat stress. The significant reduction in mutation rates and the altered mutation profiles in *atPolλ-1* plants exposed to heat stress indicate that DNA Polλ plays an important role in heat-induced mutagenesis, likely through its error-prone repair activity at damaged nucleotides that accumulate during heat stress.

### Transcriptomic analyses reveal enriched DNA Pol λ expression in central zone of meristematic cells

Studies in human cells have shown that error-prone DNA polymerases, including the DNA Polλ homolog, undergo strict spatial and temporal regulation to maintain the critical balance between genome integrity and mutagenesis ^22,23^. To better understand DNA Polλ expression patterns during plant development and growth, we explored gene expression data using the *Arabidopsis* electronic Fluorescent Pictograph (eFP) Browser ^24^. We examined DNA Polλ expression at two early developmental stages: germinating seeds and 1-week-old seedlings. In germinating seeds, DNA Polλ was expressed at low levels throughout the plants, with slightly higher expression in roots (0.9 ± 0.02) compared to shoots (0.43 ± 0.21 in the hypocotyl; 0.23 ± 0.22 in the cotyledons and vegetative shoot apical meristem (SAM)). Within the vegetative SAM, DNA Polλ showed stronger expression in central meristematic regions compared to leaf primordia (Fig. 5A). In 1-week-old seedlings, while DNA Polλ transcript levels remained generally low in the SAM and leaves, they were relatively enriched in meristematic and dividing cells of leaf primordia compared to mature leaf tissues. Notably, within the vegetative SAM, the highest expression levels were observed in the central zones (Fig. 5A).

**Figure 5:**
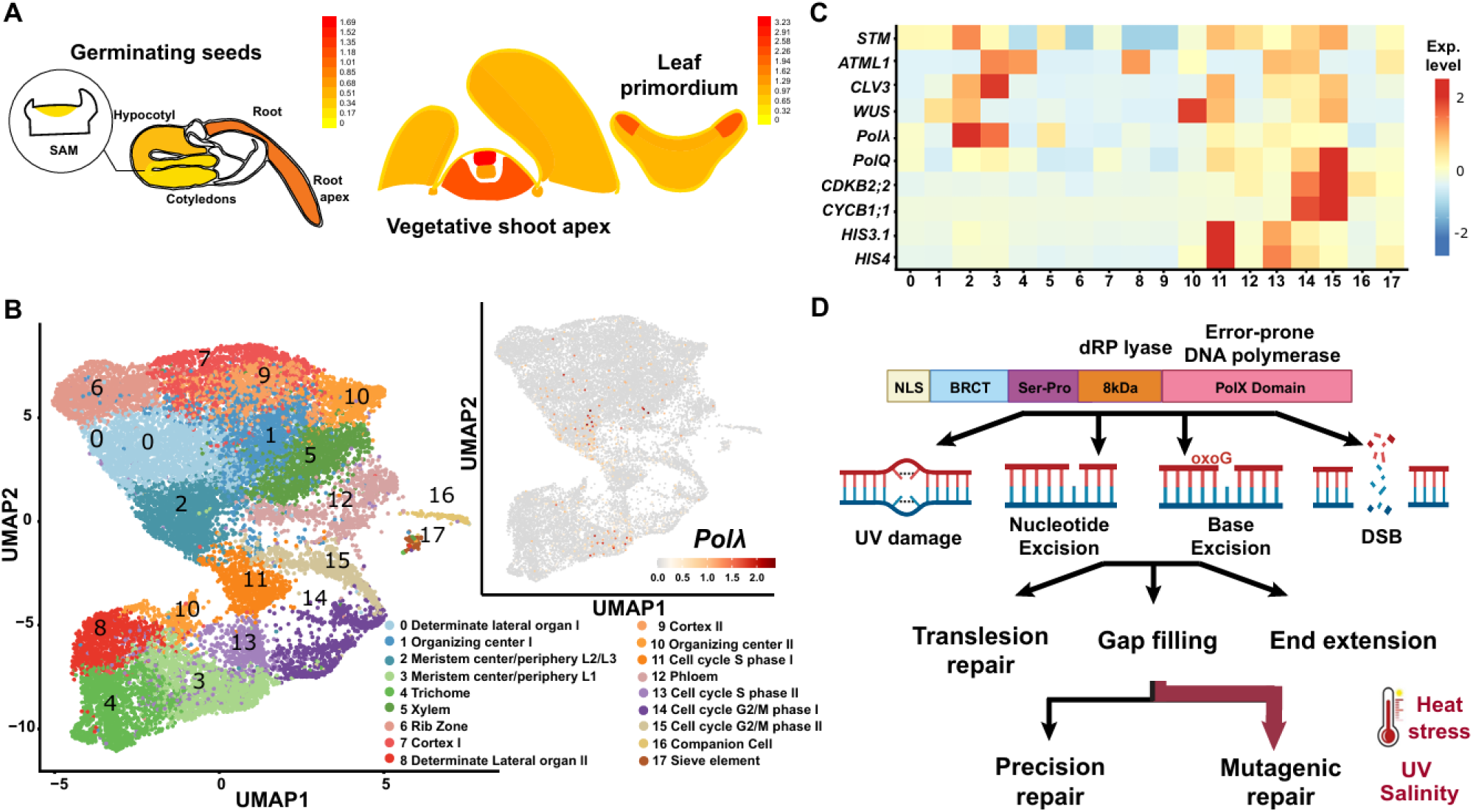
DNA Pol λ expression pattern revealed through transcriptomic analyses. **a**, Analysis of DNA Polλ expression patterns using the Arabidopsis electronic Fluorescent Pictograph (eFP) Browser in germinating seeds and 1-week-old seedlings. **b**, UMAP visualization of single-cell RNA-seq from the *Arabidopsis* inflorescence SAM (left) with DNA Polλ expression levels shown on the right. **c**, Heatmap analysis of DNA Polλ expression across cell type clusters. The highest expression of DNA Polλ is observed in the central zone of meristematic cells of the internal non-epidermal layers (L2 and L3), followed by the L1 epidermal layer. Expression patterns of the key marker genes and a specialized DNA polymerase, PolQ, for each cell type cluster are also included in heatmap. **d**, Proposed model of DNA Polλ’s role in DNA repair, maintenance of genomic stability and stress-induced mutagenesis with the versatile nature of DNA Polλ.

To gain further insight into DNA Polλ expression patterns at reproductive stages, we investigated DNA Polλ in the single-cell RNA-seq data of the inflorescence SAM of *Arabidopsis thaliana* (Fig. 5B and C) ^25^. This atlas includes transcriptomic profiles of thousands of individual cells annotated by tissue layer, developmental zone, and transcriptional identity. We specifically interrogated DNA Polλ expression across these annotated clusters and found that it was most highly expressed in central zone meristematic cells of the internal non-epidermal layers (L2 and L3; Cluster 2 in Fig. 5B and C), with the second-highest levels observed in cells from the L1 epidermal layer (Cluster 3 in Fig. 5B and C). This expression pattern in reproductive tissues parallels our observations in vegetative tissues from young seedlings. For comparison, we examined expression patterns of DNA Polymerase Q (Pol Q), a specialized polymerase mediating microhomology-mediated end joining repair (MMEJ). In contrast to the tissue-specific expression pattern of DNA Polλ, Pol Q demonstrated cell cycle-associated expression, with enrichment primarily in cells at G2/M phases rather than in specific cell types (Fig. 5C and Supplemental Fig. 5). Taken together, these transcriptomic analyses reveal that DNA Polλ exhibits distinct regulation with preferential expression in central zone meristematic cells throughout development, suggesting a specialized role in maintaining genomic integrity in these critical stem cell populations.

## Discussion

In this study, we sought to address how heat stress affects DNA repair and induces mutations both locally and globally using *Arabidopsis* as the model plant. At the local level, our findings demonstrate that heat stress significantly increases CRISPR-Cas9 targeted mutation efficiency across all chromatin contexts, with particularly dramatic effects in heterochromatic regions, showing up to 29.9-fold increases in traditionally difficult-to-edit sequences. Moreover, heat stress shifts mutation outcomes toward 1-bp insertions, regardless of chromatin states. We subsequently identified a heat-inducible error-prone DNA Polymerase, DNA Polλ, as the mediator of this shift in mutation profiles. When extending our investigation from targeted mutations to genome-wide effects using whole-genome sequencing (WGS), our data indicate that increases in genomic mutation rates under heat stress are dependent on DNA Polλ.

Our observation that heat stress enhances CRISPR-Cas9 induced mutation efficiency is consistent with previous studies ^12,14^. Yet our work further demonstrates that this enhancement occurs across diverse chromatin contexts, with the most pronounced effects in heterochromatic regions. This is particularly significant as heterochromatin has traditionally been a barrier to efficient genome editing, often requiring specialized approaches or chromatin-modifying agents to achieve reasonable editing rates ^13^. Thus, heat stress can serve as a simple and effective strategy to overcome this limitation. The molecular basis for this enhancement could involve multiple factors including increased Cas9 activity, changes in chromatin accessibility, and alterations in DNA repair mechanisms. While further research is needed to dissect their relative contributions, the differential sensitivity between euchromatic and heterochromatic regions—with heterochromatin showing more dramatic improvement at higher temperatures— points to heat-induced chromatin remodeling as one of the predominant contributors. This hypothesis is supported by previous studies demonstrating that elevated temperatures can disrupt higher-order chromatin structures in plants ^26^.

A particular intriguing finding of our study is the shift in CRISPR-Cas9 mutation outcomes toward 1-bp insertions under heat stress regardless of chromatin context or mutation rates. Notably, similar observations have been reported in analyzing CRISPR-Cas9 mutation profiles from heat-stressed rice calli ^27^, suggesting a conserved underlying mechanism across plant species. Previous research has indicated that 1-bp insertions at CRISPR-induced breaks are primarily mediated by DNA Polλ ^15,28^. Our results with the *polλ* mutant provide compelling evidence that DNA Polλ is responsible for the changes in heat-induced mutation profiles. Additionally, the 4.6-fold induction of DNA Polλ expression under heat stress offers a mechanistic explanation for the increased 1-bp insertions, establishing the links between heat exposure, DNA Polλ upregulation, and altered repair outcomes. Moreover, prior studies revealed that *Arabidopsis* DNA Pol λ physically interacted with the molecular chaperone Heat Shock Protein 90 (HSP90), suggesting an additional layer of heat-responsive regulation at the protein level^33^. Interestingly, previous research has demonstrated that *Arabidopsis* DNA Polλ expression also increases in response to other environmental stressors such as UV-B radiation and high salinity ^16,29,30^. Collectively, these observations indicated the critical role of DNA Pol λ in responding to various environmental stressors and repairing stress-induced DNA damages.

While our CRISPR-Cas9 experiments provide evidence at targeted genomic loci, our genome-wide mutagenesis analyses further demonstrate the broader role of DNA Polλ in stress-induced mutagenesis at the whole genome level. Heat stress can induce various types of DNA damage, primarily base damage caused by reactive oxygen species ^31^. DNA Polλ participates in multiple repair pathways including base excision, nucleotide excision, and translesion synthesis in addition to double stranded DNA repair. This function explains our observation of significantly reduced transversion mutations in the mutant plants under both control and heat stress conditions, consistent with the error-prone nature of DNA Polλ, particularly in bypassing oxidative DNA lesions (8-oxoG) under heat stress ^19^. Additionally, the reduction in transition mutations observed under heat stress in mutant plants suggests that DNA Polλ is also involved in promoting transition mutations during heat stress. Additional experiments are needed to fully test this hypothesis.

The error-prone nature of DNA Polλ necessitates proper regulation of its expression. The previous studies in human cells have indicated that expression of DNA Polλ is highly regulated, where overexpression of DNA Polλ is linked to higher mutagenesis in cancer cells ^32^. This aligns with our observation of tight regulation of DNA Polλ expression in plants, providing evidence for a conserved evolutionary role in modulating mutation rates across diverse species. Furthermore, our analyses revealed intriguing spatial expression patterns, with highest expression observed in meristematic regions, especially in the central zone of the shoot apical meristem. These tissue-specific patterns suggest that DNA Polλ serves multifaceted functions: maintaining genomic integrity in critical meristematic tissues while simultaneously generating stress-responsive mutations that can be transmitted to the next generation.

Taken together, we propose a model where heat stress induces increased DNA Polλ expression, resulting in higher genome-wide mutation rates (Fig. 5D). This mechanistic framework provides a molecular basis for our observed mutation profiles and highlights DNA Polλ as a key mediator in the plant’s response to elevated temperatures and other stressors, such as UV radiation and high salinity. Our model demonstrates that DNA repair genes, such as specialized DNA polymerases function not merely as guardians of genomic stability, but also as modulators of genetic diversity that facilitate adaptation to environmental stress. By elucidating the specific functions of DNA Polλ under stress conditions, our findings enhance our understanding of how plants cope with environmental stressors at the genomic level while also providing practical insights for improving genome editing efficiencies in challenging chromatin contexts.

## Methods

### Plant materials, growth conditions, and heat stress procedures

Previously described homozygous knockout lines *polλ-1* (AT1G10520; SALK_075391C) and *hos1-3* (AT2G39810; CS_844204) seeds were obtained from the *Arabidopsis* Biological Resource Center (ABRC) ^11,20^. T-DNA insertions were confirmed using primers designed with the Salk T-DNA primer design tool. Wild-type plants used in this study belong to the *Arabidopsis thaliana* ecotype Columbia-0 (Col-0). Seeds carrying the CRISPR-Cas9 T-DNA construct targeting the CHLI2 and MC4 sites were from the T2 generation and confirmed for editing as previously described ^13^.

Sterilized seeds were stratified at 4°C for 3 days, then plated on ½ Murashige Skoog (MS) media containing 0.5% sucrose and 0.8% Phytagel (Sigma-Aldrich Inc, Darmstadt, Germany). Following plating, seeds were incubated at 22°C under long day light conditions (16-hour light/8- hour dark) for 24 hours before starting the heat stress treatments. Heat stress was performed under long day conditions at 37°C for 24 hours, followed by a 24-hour recovery period at 22°C. This 48-hour cycle (stress + recovery) was repeated 3 times after which plants were allowed to recover for 2 weeks. The heat stress regimes (0x, 1x, 2x, 3x) refer to the number of times this 48-hour cycle was repeated. For temperature gradient experiments, we applied three complete cycles (3×) at 22°C, 27°C, 32°C, and 37°C. To assess heat stress responses in *polλ-1* and *hos1-3* mutants, 5-day-old seedlings were subjected to continuous heat stress at 37°C for 72 hours, followed by 3 days recovery at 22°C. Survival rates were calculated based on the proportion of non-albino seedlings relative to total seedlings. Root lengths of vertically grown seedlings were measured using ImageJ.

### Next generation amplicon sequencing

Genomic DNA was extracted using the rapid DNA extraction protocol ^33^. CHLI2 and MC4 sites were amplified using GoTaq Green Master Mix (Promega Inc. WI, USA) with an annealing temperature of 60°C and an extension time of 45 seconds; primers can be found in Supplemental Table 1. PCR amplicons were purified using QiaQuick PCR Purification kit (Qiagen, CA, USA) and sent for paired-end Next Generation Amplicon sequencing (Azenta Inc, NJ, USA). Sequencing results were analyzed using CRISPResso2 to quantify editing efficiency and repair outcomes using the default parameter reported previously ^15,34^. Three key parameters (to ignore substitutions, quantification window size 3, and minimum identity score of 90) were included to improve read selection and quantification quality.

### *Arabidopsis* floral dip transformation

Floral dip-based transformation was performed as previously described ^35^. Briefly, plants were transformed when secondary bolts were about 2-10cm with a few open flowers, approximately 5-7 days after clipping the primary bolt. All siliques were removed prior to dipping. Agrobacterium containing the T-DNA construct was cultured at 30°C for 24 hours in 50 ml for the first culture and 500ml for the second culture. Cells were harvested by centrifugation (20 minutes at room temperature at 5000 rpm) and resuspended in 500ml of MS media with 5% sucrose. Before dipping, Silwet L-77 was added to the solution at 0.02% (200ul/L). Above ground tissues were dipped into the Agrobacterium solution for 10 seconds using gentle agitation. Dipped plants were kept horizontally under dark, high-humidity conditions for 24 hours, then returned to standard growth conditions for 2-3 weeks prior to harvesting seeds.

### Quantitative RT-PCR analysis

Wild type and *atPolλ-1* seeds were grown and heat stressed as described above. After the third heat stress, seedlings were immediately collected and frozen in liquid nitrogen for RNA extraction using the RNeasy Plant Mini Kit (Qiagen, CA, USA). Reverse transcription was performed using the LunaScript RT SuperMix Kit (New England Biolabs Inc, Ipswich, MA) and no reverse transcriptase (NRT) controls were included to check for genomic contamination. The Luna® Universal qPCR Master Mix (New England Biolabs Inc, Ipswich, MA) was used for qPCR. qPCR reactions were run on a CFX Connect Real-Time System with the following parameters: 95°C for 1 minute, 38 cycles of 95°C for 15 seconds and 60°C for 30 seconds; followed by a melt curve of 95°C for 10 seconds, 65°C for 0.5 seconds, and 95°C for 5 seconds. RT-qPCR primer sequences are listed in Supplemental Table 1. EF1a was used as the housekeeping gene as it has relatively even expression across the heat stress and control ^17^. AtHSFA2 was used as the heat stress control. Relative expression levels were calculated using the ΔΔCt method (Source Data Figure 3).

### Whole genome sequencing analyses

Twelve samples (six wild-type, six *atPolλ*-1) representing three biological replicates per treatment were sent for whole genome sequencing through the NovaSeqXPlus, 150PE (Novogene, CA, USA). All heat stressed samples underwent the 3x 37°C heat stress protocol as described above. Sequencing quality information can be found in Supplemental Table 2. We followed previously described variant calling and verification methods ^21^. Read trimming was performed with trimmomatic (phred= 33) and duplicates were marked with the samtools markdup function ^36,37^. Reads were then aligned to the TAIR10 genome using BWA ^38^. Variants were called with Strelka2 in somatic TUMOR-NORMAL mode, comparing each sample with all 11 other samples as both the normal and tumor sample ^39^. Post variant calling, stringent quality filters were applied: variants had to have a reference depth of >20, with ≥ 5 reads supporting the variant. Only variants unique to each single sample were kept. Final quality checks were done by hand using IGV ^40^.

### Transcriptomic analyses

Expression patterns of DNA Polλ were analyzed using the Arabidopsis electronic Fluorescent Pictograph (eFP) Browser to assess spatial patterns in germinating seeds and 1-week-old seedlings ^24^. For reproductive stages, single-cell RNA-seq data from the Arabidopsis inflorescence shoot apical meristem were used to evaluate expression across annotated clusters defined by tissue layer and developmental zone ^25^. UMAP plots of single-cell clusters were generated using Seurat V4 (https://satijalab.org/seurat/articles/get_started.html), and gene expression analysis was visualized using either FeaturePlot (UMAP) or heatmaps. Developmental housekeeping genes included: shoot meristemless (STM; AT1G62360), meristem layer 1 (ATML1; AT4G21750), clavata3 (CLV3; AT2G27250), wuschel (WUS; AT2G17950), polymerase theta (POLQ; AT4G32700), cyclin-dependent kinase B2;2 (CDKB2;2; AT1G20930), cyclin-dependent kinase B1;1 (CDKB1;1; AT3G54180), histone 3.1 (HIS3.1;AT5G10390), histone 4 (HIS4; AT2G28740).

## Data Availability

The Data can be found at NCBI Sequence Read Archive under PRJNA1253746. All CRISPResso and WGS code can be found here: https://github.com/ZhangLab-UMN/Wootan_DNARepair_manuscript/tree/main

## Acknowledgements

C.W. is supported by Bayer Crop Science/University of Minnesota Multifunctional Agriculture Initiative Graduate Student Fellowship. F.Z. are partially supported by the National Science Foundation (IOS-2040218 and IOS-2206920) awards. X.X. acknowledges support from University of California, Davis (UC Davis) new faculty start-up funds and California Agricultural Experiment Station/National Institute of Food and Agriculture (CA-D-PLB-2850-H).

## Author Contributions

C.W. and F.Z. conceived and planned the study. C.W., J.L and X.X. performed the experiments. C.W., X.X. N.S. and F.Z. analyzed the data. C.W., X.X., N.S. and F.Z., wrote and edited the manuscript with input from all authors.

---

**Supplemental Fig 1.**
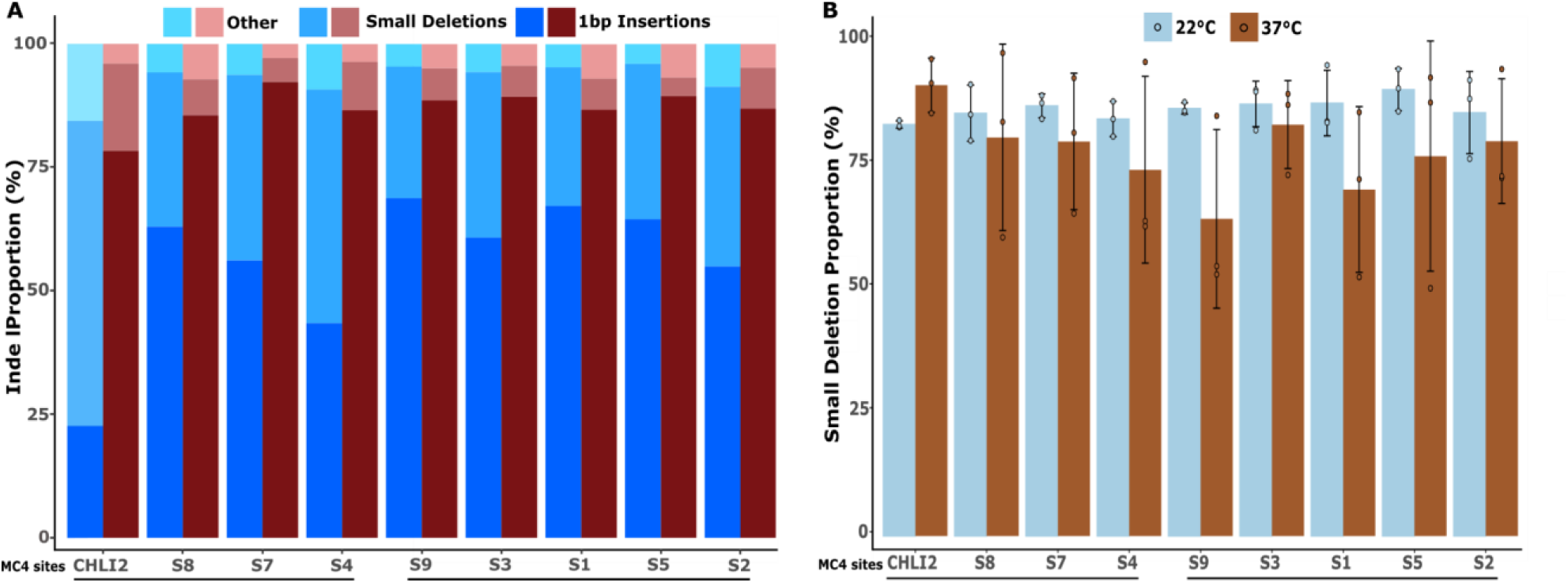
Heat stress alters 1bp insertions. **a**, All repair outcomes from control (blue hues) and heat stress (red hues) for CHLI2 and the eight MC4 sites. Indels were grouped into 1bp insertions (dark blue or dark red), 1-10 bp deletions (medium blue or medium red), or other meaning large insertions or deletions (light blue or light red). **b**, The proportion of small deletions (1-10bp) compared to all other deletions (>10 bp)) for control (light blue) and heat stress (brown).

**Supplemental Fig 2.**
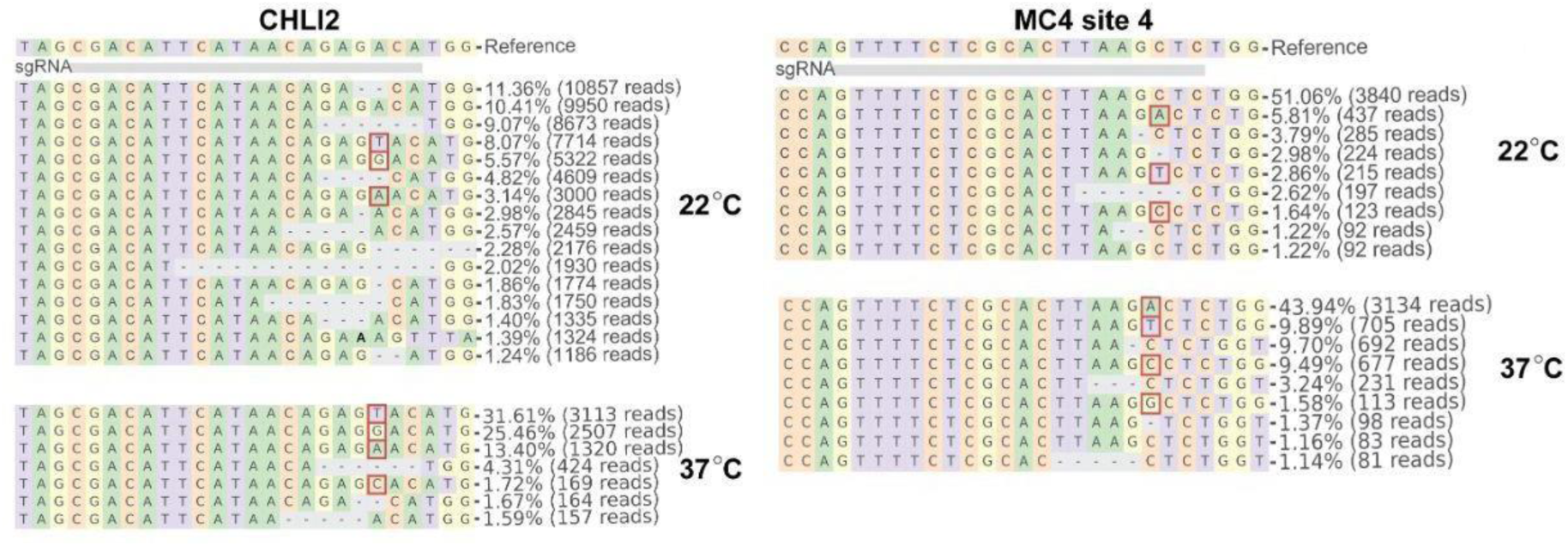
CRISPRESSO output for the CHLI2 and MC4_4 sites under control and heat stress conditions. The targeted sequences are shown on the top with 20 bp sgRNA target sites highlighted with grey bars. The 1-bp insertions are highlighted with red boxes. Small deletions are represented with dashed lines. The total read numbers and mutation rates are shown on the right.

**Supplemental Fig 3.**
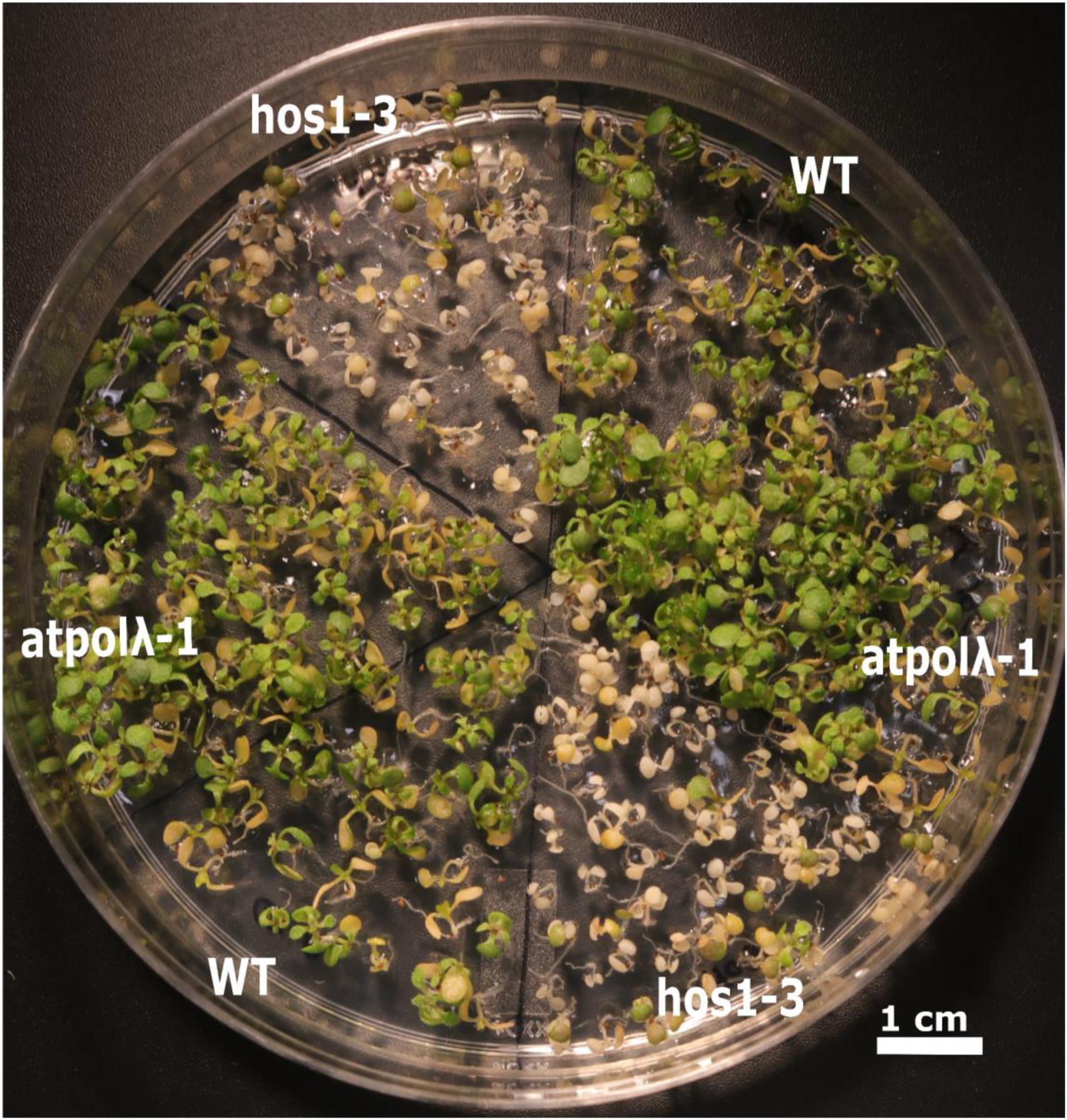
Seedling survival test under heat stress. Comparison of wild-type, hos1, and atpolλ-1 seedlings after exposure to 37°C under long day conditions for 3 days. Each plate contained two replicates of each genotype arranged diagonally. Seedlings were scored as non-viable if they exhibited severe bleaching with no post-heat stress recovery growth.

**Supplemental Fig 4.**
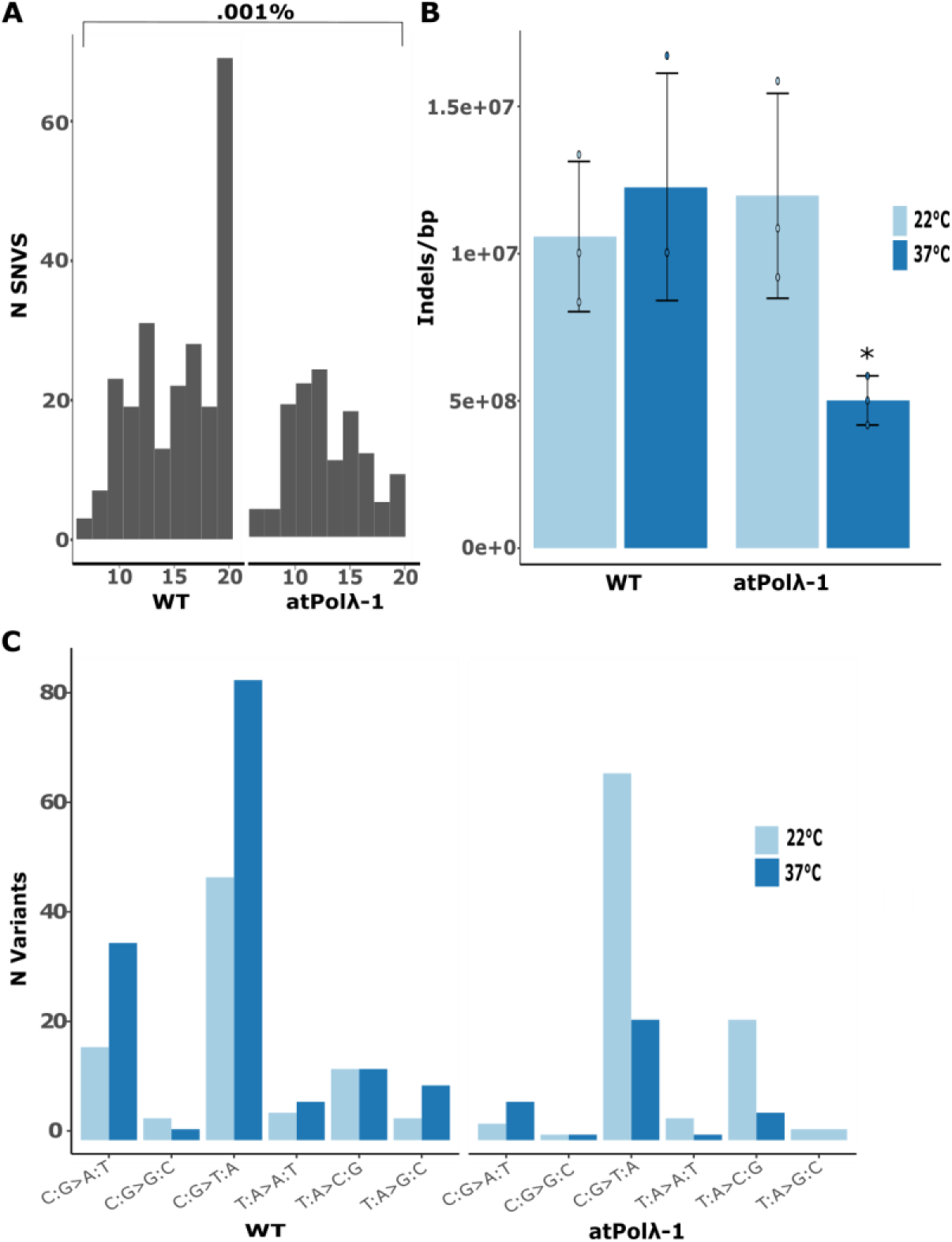
Whole genome sequencing analyses on the wild type and atpolλ-1 mutants. **a**, Empirical Variant Score (EVS) of all SNVs. All included variants have high EVS and account for 0.001% of all SNVs called. **b**, Rate of indel mutations per base pair for wild-type and atpolλ-1 under control (light blue) and heat stress (dark blue) conditions (n=3; p < 0.05). **c**, Number of mutation substitution types for atpolλ-1 and wild type under control (light blue) and heat stress (dark blue) conditions.

**Supplemental Fig 5.**
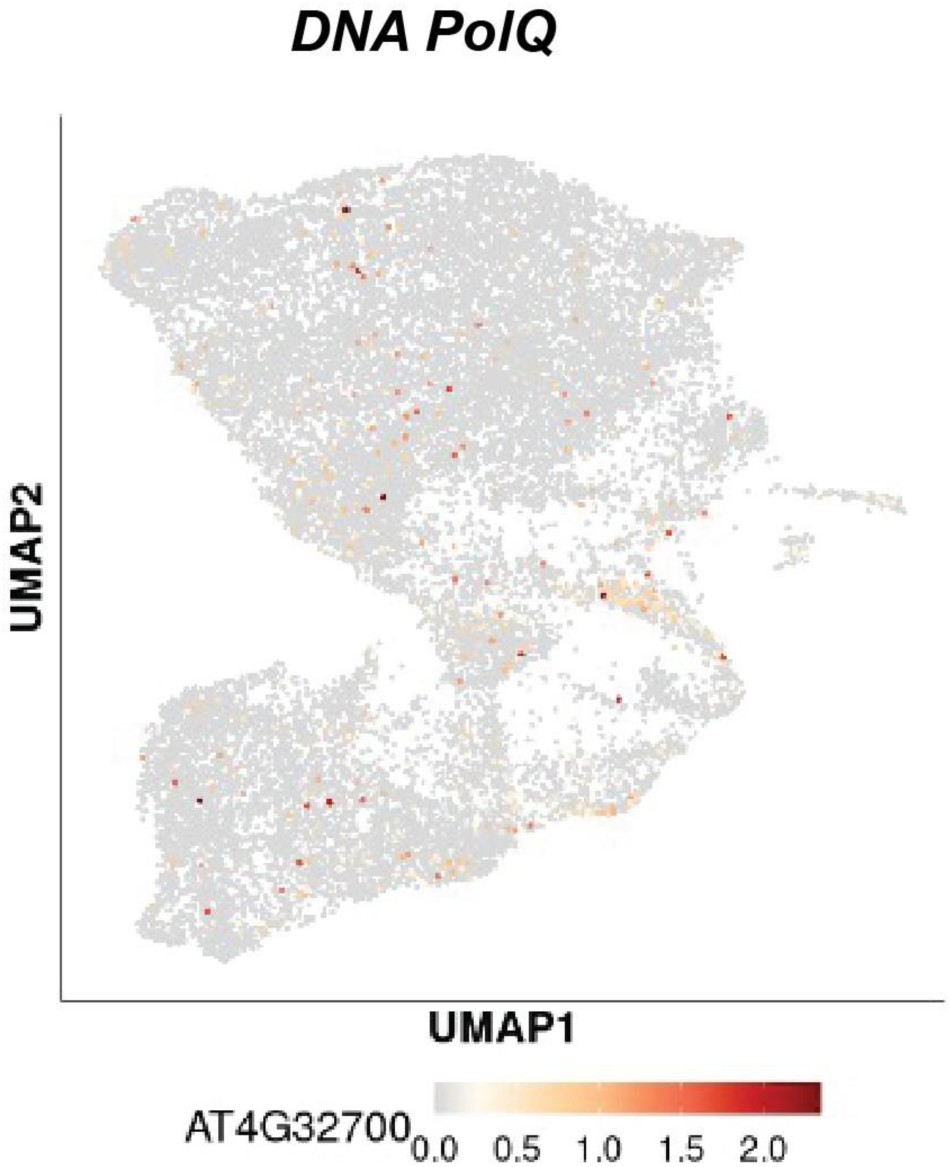
UMAP visualization of single-cell RNA-seq from the Arabidopsis inflorescence SAM with DNA PolQ (AT4G32700) expression levels.

**Supplement Table 1.**
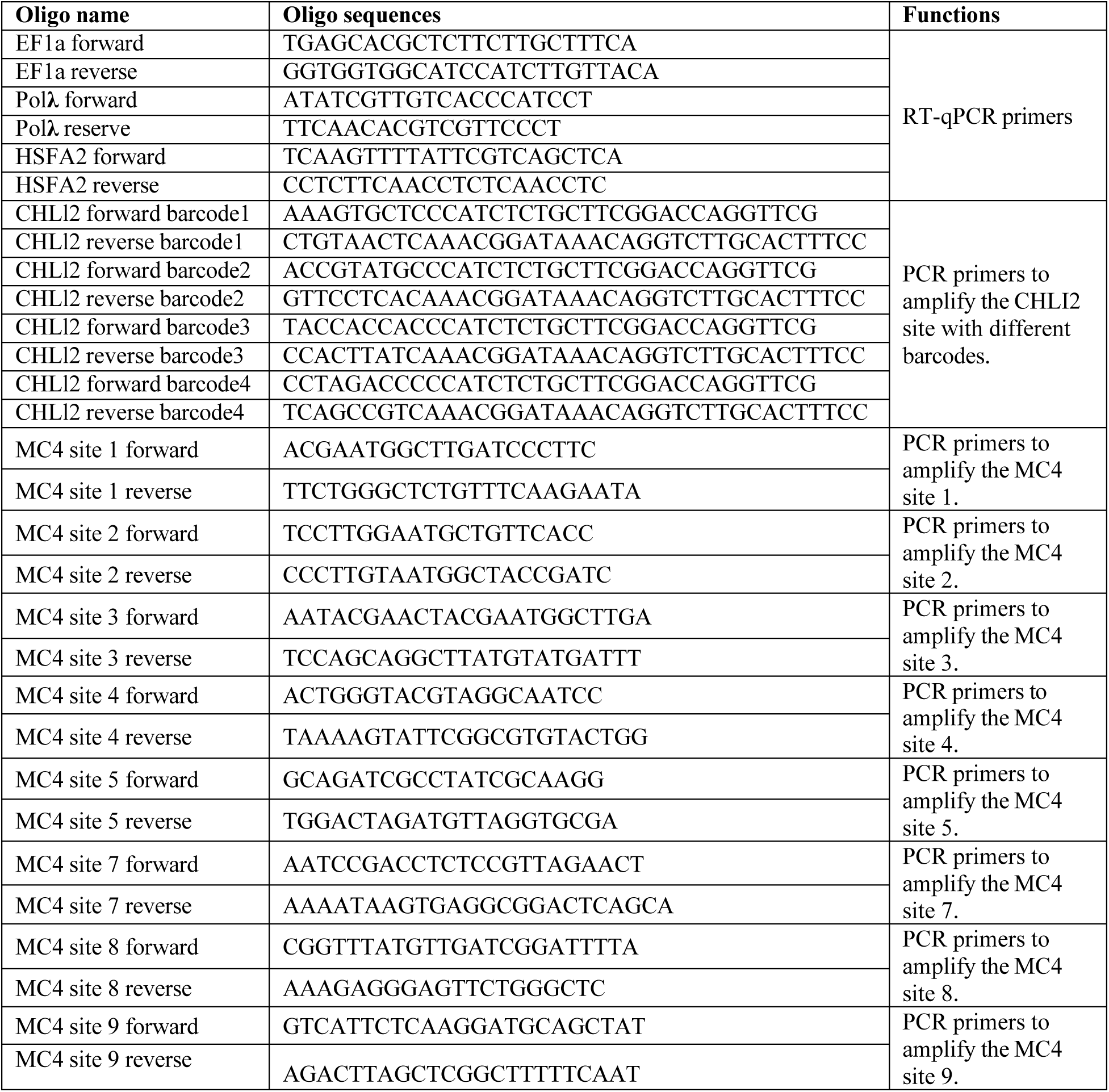
Summary of synthesized oligos.

**Supplement Table 2.**
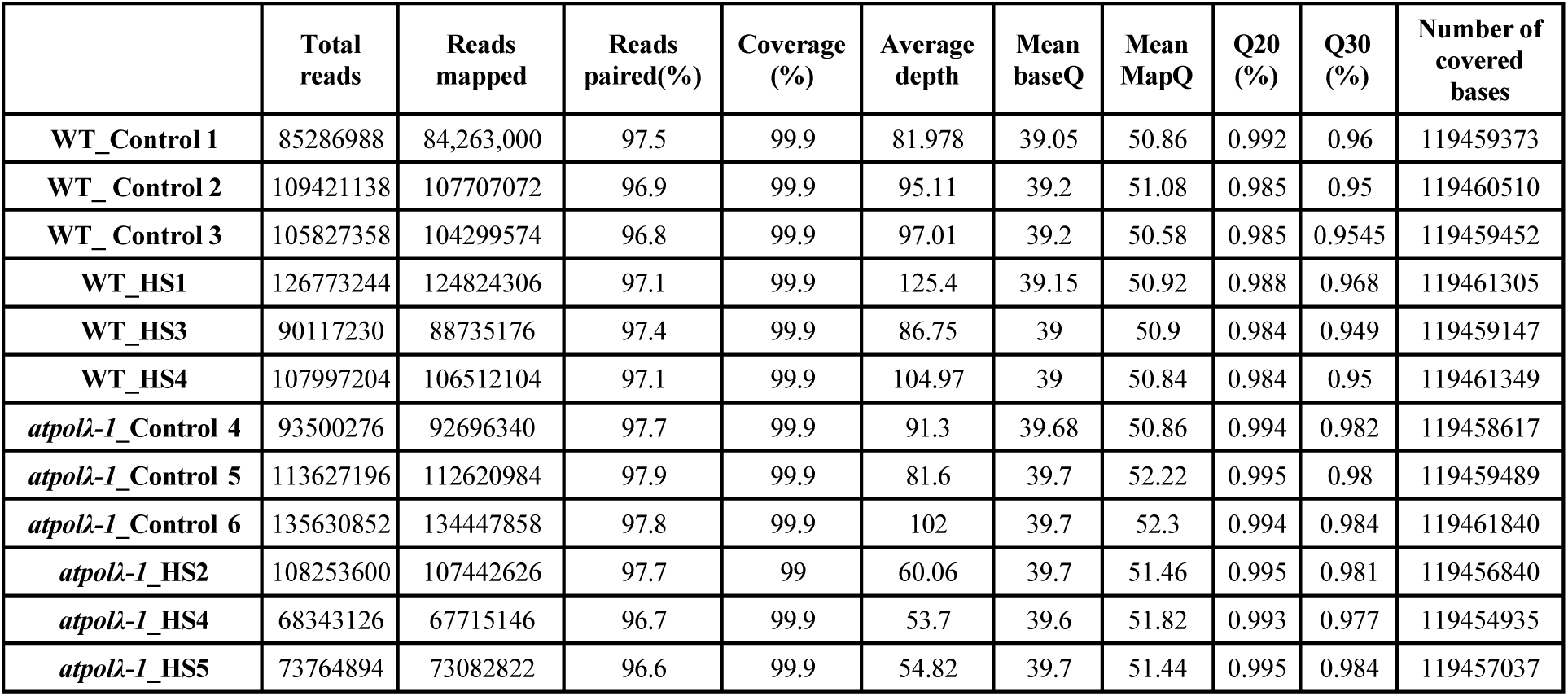
Summary of whole genome sequencing reads data.

